# Long-term reduction of short wavelength light affects sustained attention and visuospatial working memory

**DOI:** 10.1101/581314

**Authors:** Aleksandra Domagalik, Halszka Oginska, Ewa Beldzik, Magdalena Fafrowicz, Tadeusz Marek

## Abstract

The short wavelength, i.e. blue light, is crucial for non-image forming effects such as entrainment of circadian system. Moreover, many studies showed that blue light enhances alertness and performance in cognitive tasks. However, most of scientific reports in this topic is based on studies using short exposure to blue or blue-enriched light and only few focused on the effects of its reduced transmittance, especially in longer period. The latter could potentially give insight into understanding if age-related sleep problems and cognitive decline are related to less amount of blue light reaching the retina, as our lenses become more yellow with age. In this study, we investigated the effects of prolonged blocking of blue light on cognitive functioning, in particular - sustained attention and visuospatial working memory, and sleep. We used amber contact lenses reducing transmittance of blue light by approximately 90% for the period of four weeks on a group of young, healthy participants. No changes were observed for measurements related to sleep and sleep-wake rhythm. The significant effect was shown both for sustained attention and visuospatial memory, i.e. the longer blocking the blue light lasted, the greater decrease in performance was observed. Additionally, the follow-up session was conducted (approximately one week after taking off the blue-blocking lenses) and revealed that in case of sustained attention this detrimental effect of blocking BL is fully reversible. Our findings provide evidence that prolonged reduction of BL exposure directly affects human cognitive functioning regardless of sleep-related conditions.

## 1 Introduction

Humans adapted their life to 24-h light-dark cycle. As a diurnal species, we are exposed to light which is necessary not only for vision, but also constitutes a powerful modulator of non-visual functions. The non-visual (or Non-Image Forming, NIF) effects of light are mediated in general by a retinal photoreceptor system built of the intrinsically photosensitive retinal ganglion cells (ipRGC, Güler et al., 2008). However, it was shown that the latter plays a key role in the regulation of the NIF effects such as circadian rhythms and pupil constriction (Czeisler et al., 1995; Güler et al., 2008; Zaidi et al., 2007) probably due to direct neuronal projections of ipRGC cells to many hypothalamic nuclei, including suprachiasmatic nuclei (master circadian pacemaker) and olivary pretectal nuclei (Hattar et al., 2006). Human and animal studies provide evidence that the NIF system detects variations in ambient irradiance and elicits a long-term modifications of circadian rhythms as well as acute changes in hormone secretion, heart rate, sleep propensity, alertness, core body temperature, retinal neurophysiology, pupillary constriction, and gene expression (reviewed by Vandewalle, Maquet, & Dijk, 2009).

In these unique ipRGC cells, the triggering signal transduction is accomplished by photopigment melanopsin, which has maximum sensitivity in the blue, i.e. short wavelength, part of the spectrum (around 480nm; Berson, Dunn, & Takao, 2002; S Hattar, Liao, Takao, Berson, & Yau, 2002). Behavioral, biochemical and neuroimaging studies, as well as subjective measurements, demonstrated that the sensitivity of the human circadian rhythm and alerting and cognitive responses to light is blue-shifted relative to the three-cone visual photopic system, thus related to melanopsin phototransduction (see reviews by Cajochen, 2007; Chellappa et al., 2011; Vandewalle et al., 2009).

Here, we focus on alerting and cognitive functions, and how these are affected by changed light condition. Studies demonstrated that exposure to monochromatic blue light improves performance, i.e. participants respond faster and with better accuracy, and at the same time reduces sleepiness in terms of subjective ratings (Lockley et al., 2006) or stimulates higher cognitive brain activity, independently of vision (Vandewalle et al., 2013). Recently, Alkozei and co-workers (2016) have shown that exposure to blue versus amber (placebo) light led to better performance on a working memory task and increased functional brain responses related to the memory process. Later, the same group used verbal learning test and demonstrated better subsequent memory recall in the participants exposed to blue light during memory consolidation when compared to individuals exposed to an amber light condition (Alkozei et al., 2017). Recently, another neuroimaging study revealed that melanopsin-based photoreception activates a cerebral network including frontal regions, classically involved in attention and oculomotor responses (Hung et al., 2017). A range of studies introduced blue-enriched white light and showed that it improves subjective alertness and performance (Viola et al., 2008), speeds response times (Newman et al., 2016) and is more effective in reducing subjective sleepiness and enhancing cognitive performance, specifically associated with tasks of sustained attention (Chellappa et al., 2011).

Most studies focusing on the effects of short wavelength light present the experiments with exposure to blue or blue-enriched light as a condition of interest. Also, in most of the cases they report the effect of short-lasting exposure to light (from minutes to few hours). Only a few studies reported the consequences of short-term blue light filtering intervention using the short-wavelength attenuated polychromatic white light or blue-light blocking glasses. It was shown that this reduction leads to decrement of performance, subjective vigilance and efficiency, and affects physiological parameters linked to sleepiness and vigilance (van de Werken et al., 2013) as well as attenuation of LED-induced melatonin suppression in the evening and decreased vigilant attention and subjective alertness before bedtime (van der Lely et al., 2015). A chronic reduction of short-wavelength was introduced by Giménez and co-workers (2014) in the study on melatonin and sleep patterns. They used soft orange contact lenses (reducing about 50% in short wavelength range) for two weeks in order to mimic, to a certain extent, the aging effects of the lens yellowing in healthy young participants. No differences were found in the melatonin measures and the effects on sleep parameters were limited.

The goal of current study was to observe long-term effects of blue light filtering. In contrast to the aforementioned study by Giménez et al. (2014), we used lenses that reduce transmittance of blue light by about 90% and introduced them for 4 weeks. We focused on the alertness (as one aspect of sustained attention) defined as the ability to achieve and maintain a certain level of cognitive performance in a given task as well as on visuospatial working memory performance. Participants were tested once a week with Psychomotor Vigilance task and sequential picture location task. Actigraphy measurements as well as Pittsburgh Sleep Quality Index and Epworth Sleepiness Scale were used to control for factors related to sleep and sleep-wake rhythm. We hypothesized that our experimental condition causes a progressive deterioration of performance similar to the effect of aging-related cognitive decline, which might be to some extent related to reduced amount of blue light reaching the retina. Furthermore, to investigate whether these changes are reversible, we introduced a follow-up session one week after returning to normal light conditions.

## 2 Materials and methods

### 2.1 Participants

Forty-eight healthy participants started the experiment and were divided in two groups differing in the type of contact lenses used: BL-blocked (BLB) group and control group (CTRL). All of them had nearsightedness (myopia) and good experience with contact lenses wearing. Each participant underwent thorough ophthalmologic exam to exclude other sight problems and passed the Ishihara test for color blindness. Participants assigned to BLB group wore the amber contact lenses reducing the transmittance of BL by approximately 90% on the 24-hour basis (UltraVision, Igel RX, water content 77%, orange tint density 40%), whereas CTRLs wore the regular contact lenses (see filter properties on Fig 1.). The lenses were adjusted according to participants’ refractive error. There were 8 dropouts due to discomfort and/or eye irritation (7 of them initially assigned to BLB group). All participants were under ophthalmologist care throughout the whole experiment. Two (out of forty participants that completed the study) were excluded due to elevated level of daytime sleepiness throughout the whole experiment; they were from the CTRL group. Thus, data obtained from 38 participants underwent further analyses.

**Figure 1.**
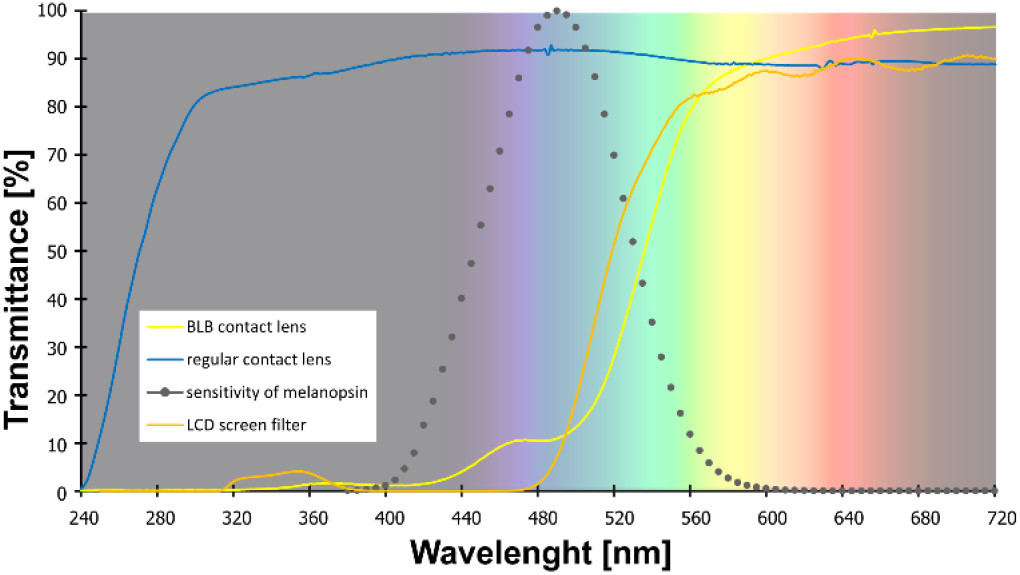
*Transmittance of the contact lenses and filter used in the study. Note: melanopsin sensitivity adapted from Irradiance Toolbox* (Lucas et al., 2014)

All participants were right-handed, with no neurological or psychiatric disorders and drug-free. During selection procedure we excluded individuals with poor sleep quality (Pittsburgh Sleep Quality Index, PSQI >6; Buysse et al., 1989) and elevated level of daytime sleepiness (Epworth Sleepiness Scale scores, ESS >10; Johns, 1991). The chronotype was assessed with Chronotype Questionnaire (Oginska et al., 2017). A T-test for independent samples was used to check the differences between groups in terms of age, sex, sleep quality, level of daytime sleepiness and chronotype (data are summarized in Table 1). No difference was found between groups. None of the participants worked night shifts or traveled across more than two time zones in the previous two months. Participants were financially rewarded for their participation, were informed about the procedure and gave their written consent. The study was approved by the bioethics commission at the Polish Military Institute of Aviation Medicine and conducted in accordance with ethical standards described in the Declaration of Helsinki.

**Table 1.** Demographic and questionnaire data.

|  | **BLB group (n=19)** | **CTRL group (n=19)** | **Difference** |
| --- | --- | --- | --- |
| **Females (nb)** | 11 | 11 | p=1.00 |
| **Age (years)** | 23.58 ± 2.76 | 24.89 ± 4.85 | p=0.31 |
| **PSQI** | 3.00 ± 1.16 | 3.00 ± 1.33 | p=1.00 |
| **ESS** | 5.90 ± 2.73 | 6.74 ± 2.05 | p=0.29 |
| **ChQ-ME** | 20.79 ± 5.63 | 20.37 ± 5.82 | p=0.82 |

*Note: Data are reported as mean ± SD (except number of females); statistical difference measured with t-test. PSQI = Pittsburgh Sleep Quality Index, ESS = Epworth Sleepiness Scale score, ChQ-ME = morningness-eveningness scale of Chronotype Questionnaire*

### 2.2 Experimental protocol

The experiment lasted six weeks (Fig 2). For each participant, the measurements were obtained once a week on the same day of the week in the evenings. Participants from CTRL group performed the task approximately at 7:30 p.m. and those from BLB group approximately at 9:30 p.m. The timing of the test differed between groups; however, it was the same in every session for each group. The goal was to match individual participants from each groups in terms of study dates, to make sure they were exposed to the same photoperiod. To check whether this discrepancy did not introduce bias regarding circadian and/or homeostatic factors, we have compared the results from the baseline session between groups. No significant differences were observed, thus the time difference between acquisition did not affect our results.

**Figure 2.**
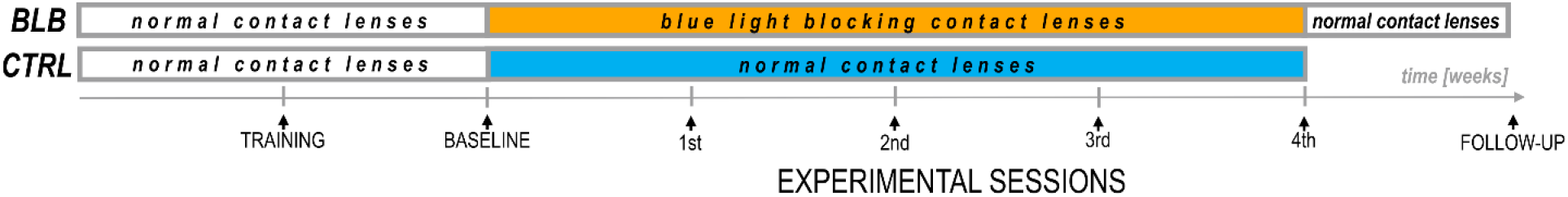
Experimental protocol.

For the first two weeks, all participants wore regular, daily disposable contact lenses. At the first session participants familiarized themselves with experimental procedures and the laboratory. The second session was treated as a baseline. For the next four weeks participants wore monthly disposable contact lenses with different filter properties according to group. Additionally, participants from BLB group had one more session (follow-up) about one week after the main experiment.

There were no behavioral restrictions imposed on participants. The experiment was conducted during spring to fall months ensuring greater availability of sunlight.

At each session, participants performed two experimental tasks in front of the computer screen and filled in the ESS questionnaire. During baseline session and on last experimental session participants additionally fulfilled the PSQI questionnaire. Through the whole experiment (six weeks) they wore actigraphs (AMI – Ambulatory Monitoring Inc. or MotionWatch8 – CamNtech Ltd.) on their nondominant wrist.

To avoid any expectancy effect, a possible positive or negative influence of the blue light filtering on functioning and the general well-being was not suggested to participants before the start of experiment. They were informed that the *research goal* is primarily, the observation of cognitive functioning in the situation of “sharpening the eye” (amber lenses were advertised as an aid for the eye to support sharp vision, especially at dusk, which can be useful for drivers and athletes).

### 2.3 Experimental tasks

Participants performed a Psychomotor Vigilance Task (PVT), a widely used test of sustained attention (Basner and Dinges, 2011; Dinges and Powell, 1985). The task required pressing a response button (with index finger) as soon as the stimulus appears, which stops the stimulus counter and displays the reaction time (RT) in milliseconds for a 1s period. It was emphasized to the participants not to press the button in the absence of stimuli, which yielded a false start warning on the screen. If a reaction was slower than 1s, the warning “too slow” appeared. The inter-trial interval varied randomly from 2 to 10s, and the task duration was 5min, comprising about 42 stimuli.

To study visuospatial working memory we used a sequential picture location task (SPLT; Fig 3). In the task, 4 pictures were sequentially presented in random square of the 4×3 grid for 500ms with 900ms interval between them. Participants were asked to remember the location of pictures. After a delay lasting from 2 to 9s, a memory probe was presented at the screen until response was given. The instruction was to respond “yes” (index finger) if the memory probe was in the same location as during the preceding sequence or “no” (middle finger) if the location was different. A 5s blank screen was presented between trials. There were 56 trials what resulted in approximately 15 min of task duration (actual task duration was dependent on participants’ response time). The pictures for the task were taken from BOSS database - the Bank of Standardized Stimuli (Brodeur et al., 2010). Participants did not perform the task during follow-up session. One participant from CTRL group was excluded from the analysis due to low performance (less than 70% accuracy in all sessions). Nine of thirty seven participants performed shorter version of task, i.e. there were 31 trials and 3 pictures to remember in each trial.

**Figure 3.**
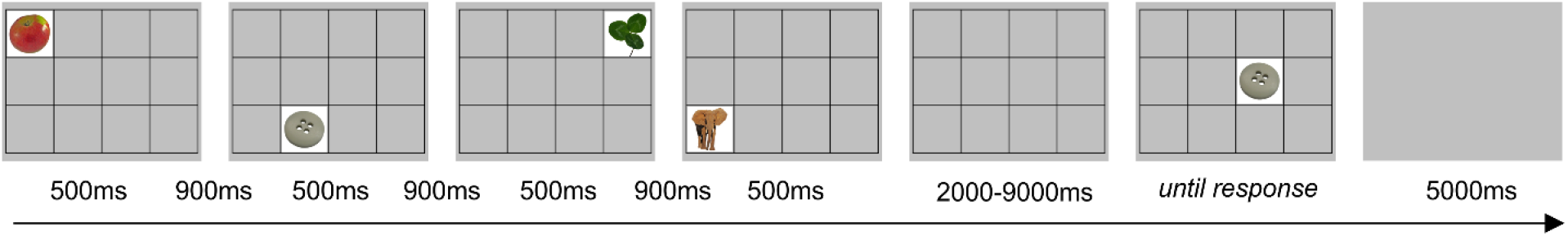
Example of trial for the SPLT task. The item shown would require a response “no”

Both tasks were performed on computer with 19-inch LCD screen. For training and baseline session for BLB group as well as for all experiment for CTRL group the blue-blocking filter was used on the screen to ensure similar visual condition (i.a. color perception) for all participants (see filter properties on Fig 1). Participants responded with arrow keys on the keyboard.

### 2.4 Data analysis

In the PVT, errors of commission were defined as responses without a stimulus or those with RT < 100ms, whereas errors of omission as lack of response on stimulus or responses with RT ≥ 500ms. The number of errors was calculated as a percentage of all stimuli in the task. For correct responses mean RT, mean RT for 10% of the fastest responses and mean RT for 10% of the slowest responses were calculated as the most frequently reported PVT outcome metrics (Basner and Dinges, 2011). First three stimuli were discarded from analysis in each PVT trial. In the SPLT task, only accuracy (in %) was measured since there was no time restriction on response.

The outcomes of both tasks were established for each experimental session and then compared using mixed analysis of variance (ANOVA) with *sessions* (5 levels accounting for one baseline and four sessions of wearing BLB/regular contact lenses) as within-subject and *group* (2 levels: BLP and CTRL group) as between-subject factors. To reveal if there is a changing pattern of task outcomes through consecutive sessions and if this pattern differs between groups, the interaction of session and group factors was reviewed. The Tukey HSD test was used for post-hoc comparisons. For the SPLT task, additional between-subject factor was added to the ANOVA analysis in order to take into account the two version of task (note: none of the effect including *task version* factor was significant).

For PVT task, indicators of state instability, i.e. standard deviation of RT and the time-on-task effect was tested and compared between sessions. The mean RT for each minute of task was calculated and introduced in mixed effect ANOVA with *minutes, sessions* and *group* as factors. Additionally, to evaluate if there is a difference in the time-on-task effect across experimental sessions, the slope of linear regression line across RT for each minute of task was calculated and compared using mixed effect ANOVA with *sessions* and *group* as factors. The standard deviation of RT was compared in the same way.

Furthermore, the comparison of the follow-up session with baseline and last week of wearing the BLB contact lenses was performed on data from BLB group with repeated measures ANOVA test. The goal of this analysis was to check whether any changes in behavior diminish approximately one week after taking off the blue light blocking contact lenses.

The questionnaire data was analyzed using analogical statistical tests.

## 3 Results

The PVT outcome metrics are presented on graphs in figure 4. Significant interaction effect of *session* and *group* factors was observed for mean RT (F_(4, 144)_=2.59, p<0.05, partial η^2^=0.07) and mean RT for 10% of the fastest responses (F_(4, 144)_=2.80, p<0.05, partial η^2^=0.07). The Tukey HSD post-hoc comparisons yielded a significant increase of these measures in consecutive sessions only for BLB group; detailed test results are presented on the graphs (Fig 4). Other measures did not reach the significance level: mean RT for 10% of the slowest responses (F_(4, 144)_=1.54, p=0.19), number of omission errors (F_(4, 144)_=1.37, p=0.25) and number of commission errors (F_(4, 144)_=1.51, p=0.20).

**Figure 4.**
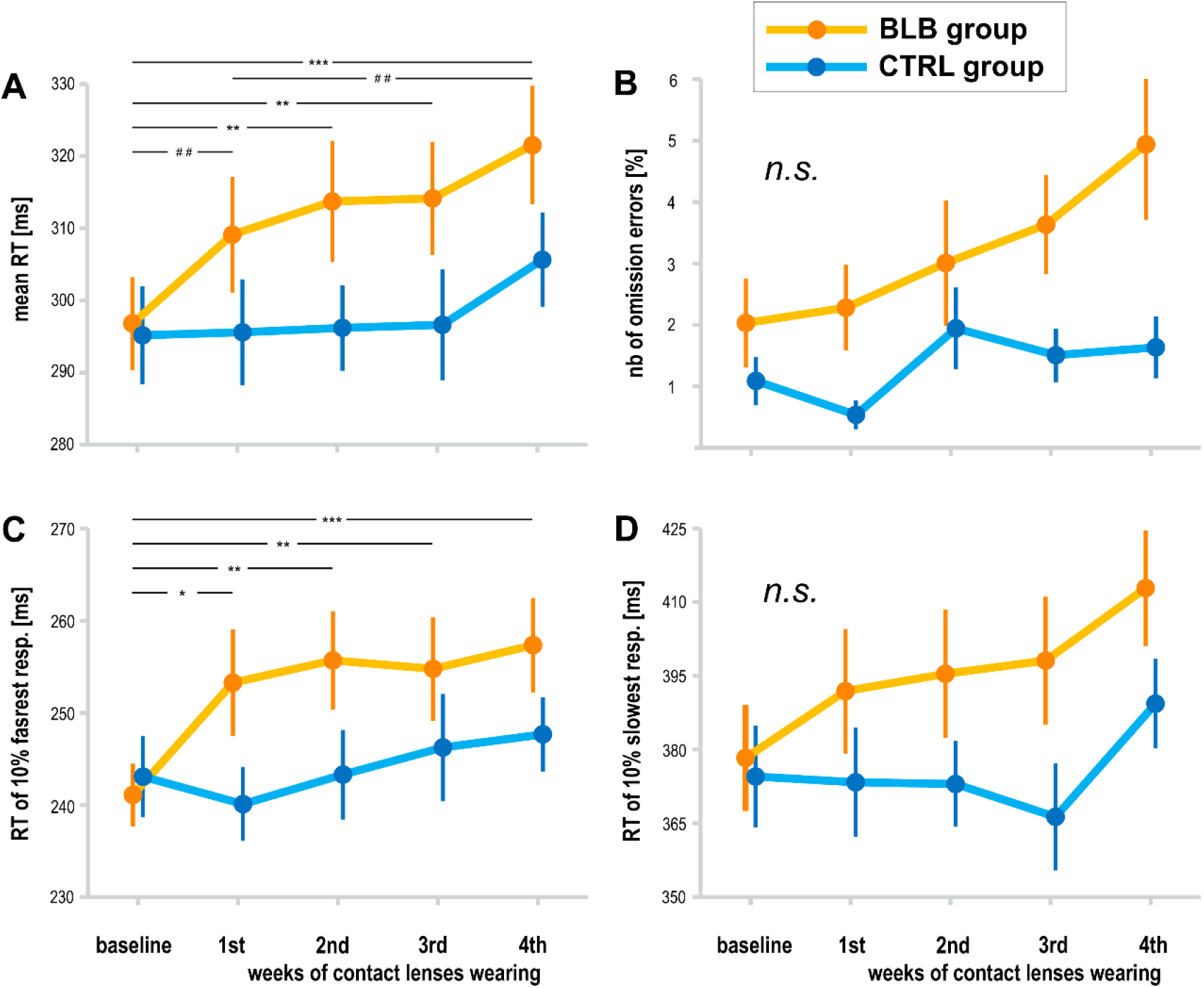
PVT outcomes for BLB and CTRL group for the baseline and 4 weeks of experimental condition: A) mean reaction time, B) number of omission errors, C) reaction time for 10% of the fastest responses, D) reaction time for 10% of the slowest responses. Note: *** denote p< 0.001; ** denote p< 0.01; * denote p< 0.05; ## denote p< 0.1; bars indicate standard error.

The time-on-task effect, i.e. increasing RT in the course of task, was present (F_(4, 144)_=9.94, p<0.001, partial η^2^=0.22), however interaction with group and session was not significant (F_(16, 576)_=1.21, p=0.25, partial η^2^=0.03). The comparison of slopes did not reveal significant interaction between sessions and groups (F_(4, 144)_=0.64, p=0.63), neither the standard deviation of RT (F_(4, 144)_=0.55, p=0.70).

For the PVT task, we compared the follow-up session with the baseline and last week of wearing the BLB contact lenses for BLB group (Fig 5). The repeated-measure ANOVA analysis revealed significant differences between those three sessions in mean RT (F_(2, 36)_=21.79, p<0.001, partial η^2^=0.55), mean RT for 10% of the fastest responses (F_(2, 36)_=12.02, p<0.001, partial η^2^=0.40), mean RT for 10% of the slowest responses (F_(2, 36)_=13.58, p<0.001, partial η^2^=0.43) and number of omission errors (F_(2, 36)_=5.15, p<0.05, partial η^2^=0.22). The post-hoc tests for all of the above mentioned analyses revealed significant difference between the baseline session and 4th week of wearing BLB contact lenses and between this session and the follow-up. There was no difference between the baseline and follow-up session. Number of commission errors did not show significant differences between sessions (F_(2, 36)_=0.13, p=0.88).

**Figure 5.**
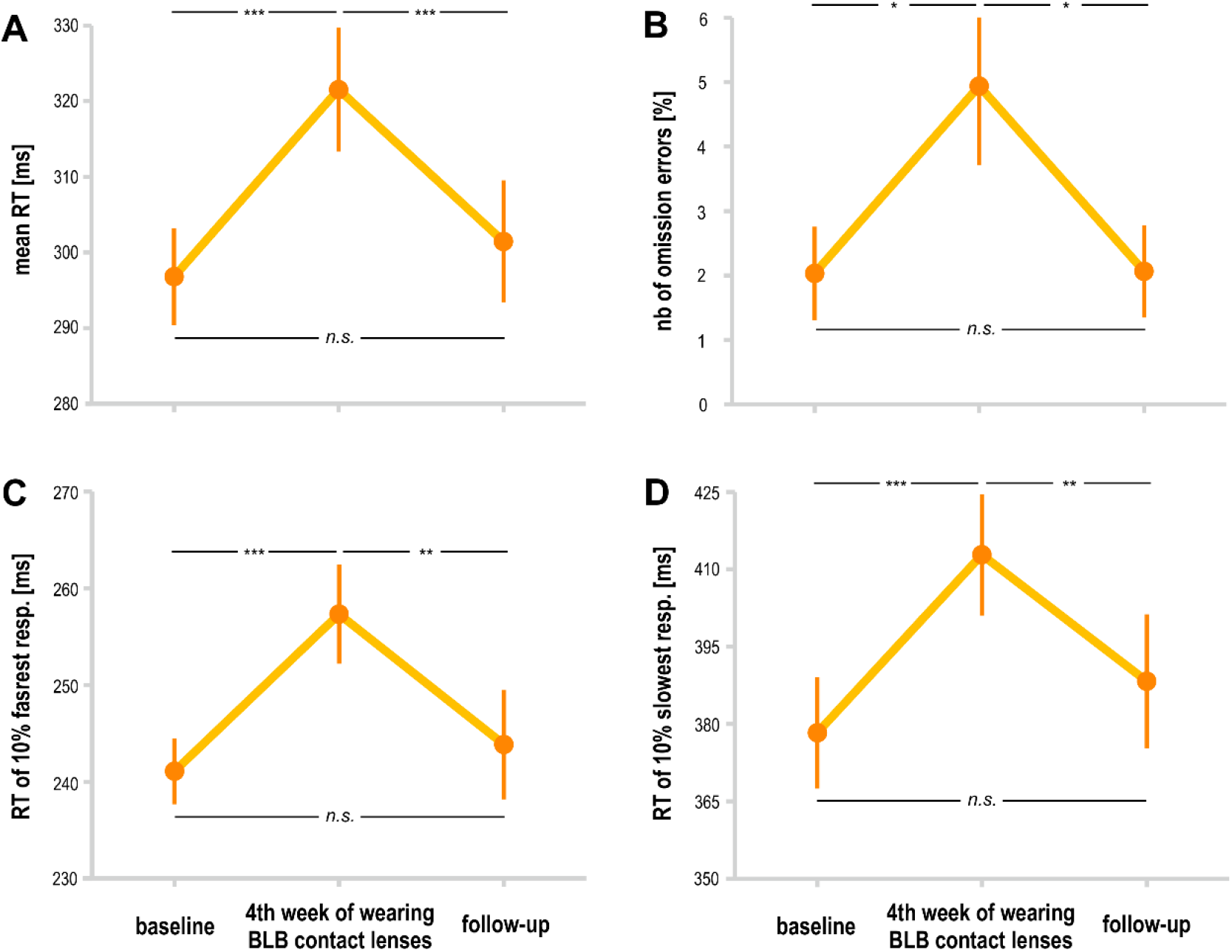
Comparison of PVT outcomes for BLB group between baseline, 4th week of blue light reduction and the follow-up session: A) mean reaction time, B) number of omission errors, C) reaction time for 10% of the fastest responses, D) reaction time for 10% of the slowest responses. Note: *** denote p< 0.001; ** denote p< 0.01; * denote p< 0.05; ## denote p< 0.1; bars indicate standard errors.

There was significant interaction between *group* and *session* for the accuracy in the SPLT task (F(4,132)=3.77, p<0.01, partial η^2^=0.10; Fig. 6). The post-hoc test revealed significant decrease in accuracy only for BLB group (see Fig 6).

**Figure 6.**
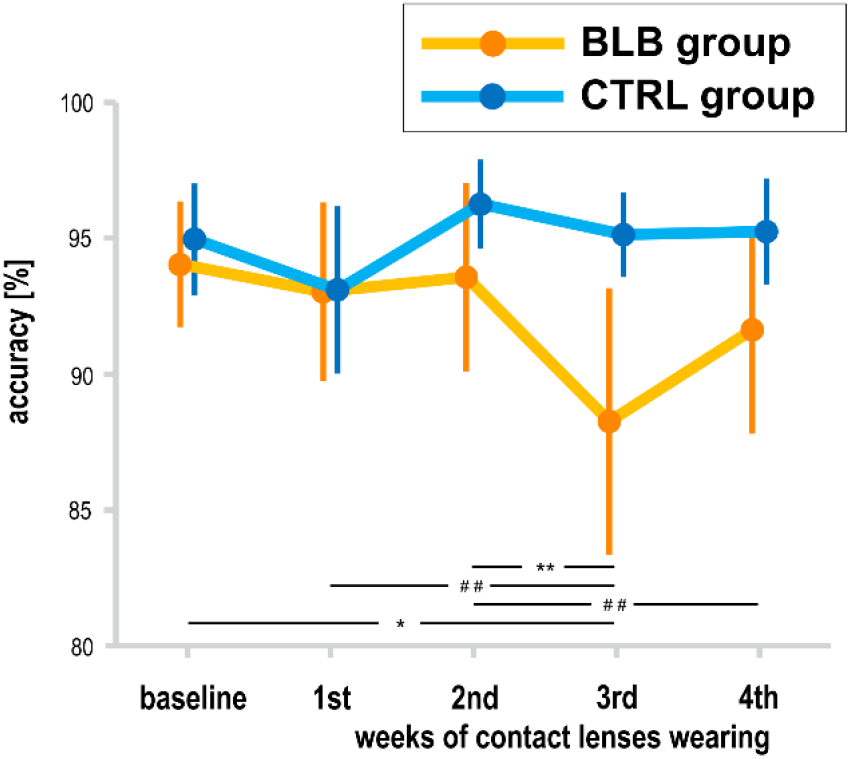
Accuracy in SPLT task for BLB and CTRL group for the baseline and 4 weeks of experimental condition. Note: *** denote p< 0.001; ** denote p< 0.01; * denote p< 0.05; ## denote p< 0.1; bars indicate standard errors.

Sleep parameters assessed with actigraphy are presented in Table 2. Actigraphy data for whole experiment from 6 participants (1 form EXP and 5 from CTRL group) were not recorded due to technical problems. There were no significant differences between groups and sessions for those measures (sleep onset: F_(4, 120)_=0.68, p=0.61; sleep offset: F_(4, 120)_=0.40, p=0.81; sleep latency: F_(4, 120)_=0.24, p=0.92; actual sleep time: F_(4, 120)_=0.72, p=0.58).

**Table 2.**
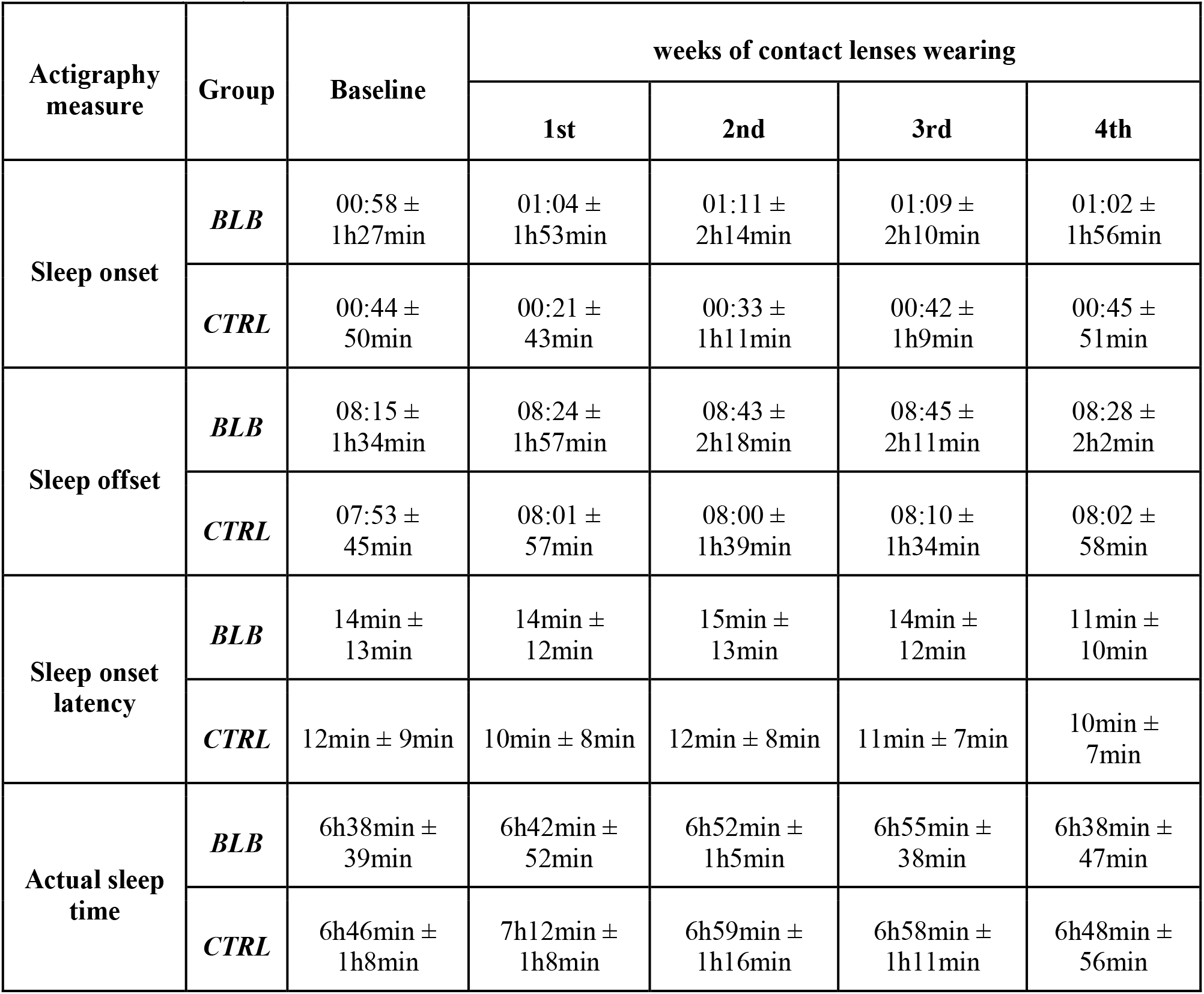
Actimetry-derived sleep parameters in baseline and during experiment period in experimental (BLB) and control groups.

| Actigraphy measure | Group | Baseline | weeks of contact lenses wearing |  |  |  |
| --- | --- | --- | --- | --- | --- | --- |
|  |  |  | 1st | 2nd | 3rd | 4th |
| Sleep onset | *BLB* | 00:58 ± 1h27min | 01:04 ± 1h53min | 01:11 ± 2h14min | 01:09 ± 2h10min | 01:02 ± 1h56min |
|  | *CTRL* | 00:44 ± 50min | 00:21 ± 43min | 00:33 ± 1h11min | 00:42 ± 1h9min | 00:45 ± 51min |
| Sleep offset | *BLB* | 08:15 ± 1h34min | 08:24 ± 1h57min | 08:43 ± 2h18min | 08:45 ± 2h11min | 08:28 ± 2h2min |
|  | *CTRL* | 07:53 ± 45min | 08:01 ± 57min | 08:00 ± 1h39min | 08:10 ± 1h34min | 08:02 ± 58min |
| Sleep onset latency | *BLB* | 14min ± 13min | 14min ± 12min | 15min ± 13min | 14min ± 12min | 11min ± 10min |
|  | *CTRL* | 12min ± 9min | 10min ± 8min | 12min ± 8min | 11min ± 7min | 10min ± 7min |
| Actual sleep time | *BLB* | 6h38min ± 39min | 6h42min ± 52min | 6h52min ± 1h5min | 6h55min ± 38min | 6h38min ± 47min |
|  | *CTRL* | 6h46min ± 1h8min | 7h12min ± 1h8min | 6h59min ± 1h16min | 6h58min ± 1h11min | 6h48min ± 56min |

The subjective sleepiness (ESS) and sleep quality (PSQI) results are presented in figure 7. The interaction effect of *session* and *group* factors in ANOVA test for both scores was not significant (ESS: F_(4, 144)_=1.72, p=0.15, PSQI: Current effect: F_(1, 36)_=0.12, p=0.73). When considering the follow-up session, ESS score did not show significant effect of session (F_(2, 36)_=2.24, p=0.12).

**Figure 7.**
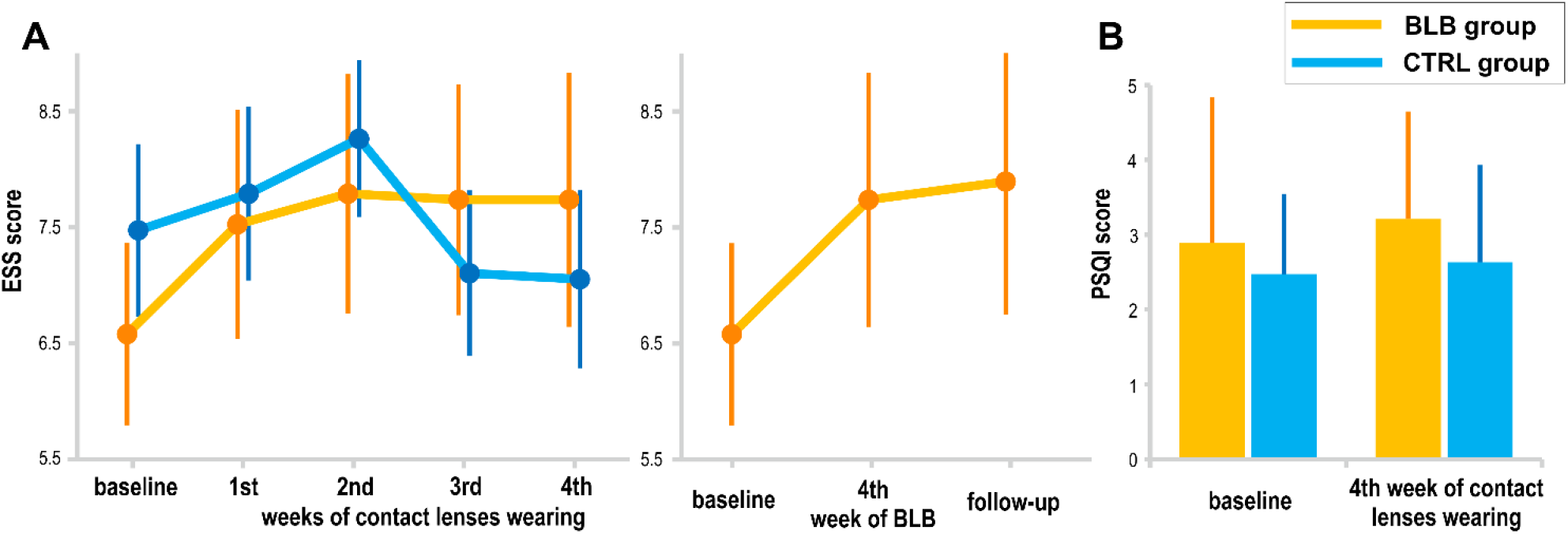
Subjective measures of sleepiness and sleep quality: A) EES score for BLB and CTRL group in consecutive experimental session as well as comparison to follow-up session for BLB group, B) PSQI score at the baseline session and after 4 weeks of experimental condition for both groups. Note: bars indicate standard errors.

## 4 Discussion

In this study we investigated the effects of prolonged blocking of blue light on sustained attention and visuospatial working memory. We used the PVT, a simple reaction time task that indicates the ability to achieve and maintain a certain level of cognitive performance (Souman et al., 2018), as well as a SPLT task requiring memory both for object and spatial locations. Actigraphy was applied to control for timing and duration of sleep, PSQI for assessing its quality, and ESS to control for daytime sleepiness levels. We compared the data collected from two groups of participants wearing either contact lenses with filter blocking approximately 90% of blue light or normal contact lenses during four consecutive weeks. Results show significant change of pattern in majority of the PVT outcome metrics in the BLB group. Particularly, the longer blocking the blue light lasted, the slower were responses as indicated by mean RT and mean of the 10% fastest responses. The effects on other PVT measures did not reach significance level, although they show also increasing pattern in the course of weeks with reduced blue light. In the CTRL groups, all the metrics were stable through all sessions. There was no effect of changed light condition on RT variability measured by standard deviation as well as on time-on-task effect. The comparison between baseline session and last experimental session in the BLB group revealed significant increase in all of the PVT outcome and significant decrease of those measures at the follow-up session. In case of the SPLT task, significant decrease in accuracy was shown only for BLB group. Sleep timing and length, subjective sleep quality and daytime sleepiness did not change during the experimental intervention.

As hypothesized, our study show that blocking blue light slows down reactions in sustained attention task and cause deterioration of performance in visuospatial memory task. This effect gradually strengthens during the consecutive weeks. Interestingly, our results revealed that in case of sustained attention this effect is reversible, as the reaction times decrease to the baseline level after returning to ‘normal’ light conditions (i.e. with BL in the spectrum).

The effect of blue light on cognitive processes may be considered in terms of complex interactions between circadian, sleep, and arousal factors. Particularly, blue light, through NIF system, can either act directly on neuronal system and thus on alertness and behavior, or indirect effects may occur due to disrupted entrainment of circadian system and/or sleep (Fisk et al., 2018). Therefore, the effects of changed blue light condition can be interpreted in terms of different mechanisms.

Taking into consideration studies on sleep-wake regulation, it can be stated that blocking blue light causes similar outcomes to those of sleep deprivation. Thus, performance decrement reported in our study might be linked to disruption of sleep or sleep-wake rhythm. However, in contrast to research on sleep deprivation, our measurements imply that participants had no sleep problems in conditions of blue light filtering. Both, sleep duration and sleep quality were not affected. Taking into account the direct impact of light on melatonin secretion and circadian regulation, one could expect, in conditions of blue light blockade, the following changes in participants’ sleep-wake pattern: earlier bedtime, shorter sleep latency, and longer sleep. This could lead to improved well-being due to better sleep. However, it may be also the case that lack of triggering effect of blue light would be the key factor of daytime drowsiness, hence - lack of energy and decreased well-being. In our study, 8 out of 18 participants (44%) showed earlier sleep onset after four weeks of experiment (so did 5 out of 14 controls, i.e. 36%), 35% of experiment participants (vs. 43% of controls) exhibited shortened sleep latency, and 39% of participants (vs. 36% of controls) had longer sleep. Those results do not support the above assumptions on direct link between blue light and sleep/wake rhythm. In general, the prolonged blue light reduction did not result in significant changes of sleep pattern, although we are aware of the fact that the ‘pristine’ internal sleep timing is influenced by external social factors that are not controlled in case of an experiment conducted in ‘natural’ conditions. Moreover, the results of Giménez et al. (2014) where blue light exposure was reduced for two weeks did not show differences in the timing of sleep, its efficiency and subjective quality, likewise no effect was found on dim light melatonin onset or on amplitude of melatonin rhythms. Thus, the unchanged sleep related indicators in our study may be interpreted as speaking for lack of BLB effect on sleep-wake rhythm and sleep *per se* and in consequence on performance through this indirect pathway.

Therefore, the results of our study could be interpreted in terms of direct effect of light on alertness. The series of neuroimaging experiments by Vandewalle and colleagues demonstrated that blue light induces modulations of brain activity while participants are engaged in non-visual cognitive tasks (Vandewalle et al., 2007a, 2007b, 2009). Those activations regarded alertness-related subcortical structures such as brainstem, hypothalamus, dorsal and posterior parts of thalamus, hippocampus and amygdala. The modulations were detected also in the cortex, in areas involved in bottom-up reorientation of attention and in the regions linked with top-down regulation of attention. At the behavioral level, blue enriched light was shown to enhance subjective alertness and led to significantly faster reaction times in tasks associated with sustained attention and working memory (Alkozei et al., 2016; Chellappa et al., 2011a). Our experimental condition is opposite, as we reduce blue light exposure and so are our results as we observe increasing reaction times and decreasing accuracy. Hence, the reduced amount of blue light reaching the retina may be associated with insufficient stimulation of the alerting and/or orienting system in the brain that in consequence have impact on cognitive processes.

Prolonged filtering of the blue light takes place in the aging process of the eye - the natural lens becomes more yellow because of accumulation of chromophores that decrease transmission especially in the short wavelength range of visible spectrum (Giménez et al., 2010; Kessel et al., 2010). It was shown the transmission of light at 480nm decreases by 72% from the age of 10 to 80 years (Kessel et al., 2010). As known, aging is often associated with sleep and circadian disturbances (Duffy and Czeisler, 2002) as well as cognitive decline (Cabeza et al.). A large sample study show that the age-related lens yellowing may be responsible for sleep problems in the elderly because of disturbed photoentrainment of circadian rhythms (Kessel et al., 2011). Our results suggest that reduced amount of blue light reaching the retina might be one of factors influencing the deterioration of cognitive functioning. A finding by Schmoll and co-workers (2011) supports this explanation.

They examined patients after cataract surgery and revealed that improved blue light transmission has a beneficial effect on cognitive function (responses became both quicker and more consistent following surgery). Moreover, the detrimental effect of blocking blue light can be reversed as demonstrated in our study (returning to baseline level of task performance after returning to normal light conditions) or attenuated as shown in studies on elderly population (a long-term, whole-day bright light exposure in large cohort of care facilities residents, (Riemersma-van der Lek et al., 2008)).

In summary, we studied the effects of prolonged reduction of blue light on sustained attention and visuospatial memory in natural conditions with simultaneous control of measures related to sleep and sleep-wake cycle. The results revealed worsening of performance during changed light conditions in comparison to control group where no filter was applied. No effects on sleep parameters were found.

Our findings demonstrate that filtering blue light affects human cognitive functioning. The lack of significant changes in sleep and sleep-wake indicators during four-week experiment may be an indirect proof of stability of sleep-wake rhythm. Thus, the observed deterioration of cognitive functioning through the decrease in ‘psychomotor vigilance’ and visuospatial working memory may be directly attributed to weakening of alerting effect of blue light.

## HIGHLIGHTS

The effects of prolonged reduction of blue light on human cognition were studied.

Filtering of blue light affects sustained attention and visuospatial memory.

The longer blocking the blue light lasted, the worse performance was observed.

No effects on sleep and sleep-wake cycle parameters were found.

## AUTHOR CONTRIBUTIONS

AD, HO, EB, MF, TM: conception or design of the work; AD, HO: acquisition and analysis; AD, HO, EB, MF, TM: interpretation of the results; AD: drafting the manuscript; AD, HO, EB, MF, TM: revising the manuscript critically and final approval of the version to be published;

## ACKNOWLEDGMENTS

The authors thank Justyna Janik for support in data acquisition, Mariusz Duda from Department of Biophysics, Faculty of Biochemistry, Biophysics and Biotechnology at the Jagiellonian University for transmittance measurements of contact lenses used in the study and Piotr Chaniecki, Agnieszka Ścisłowicz, Jagoda Miszczyk, Paulina Renke from Ophthalmology Clinic at 5th Military Clinical Hospital in Krakow for participants’ eyes examination.

